# Local population structure in Cambridgeshire during the Roman occupation

**DOI:** 10.1101/2023.07.31.551265

**Authors:** Christiana L. Scheib, Ruoyun Hui, Alice K. Rose, Anu Solnik, Eugenia D’Atanasio, Sarah A. Inskip, Craig Cessford, Samuel J. Griffith, Rob Wiseman, Benjamin Neil, Trish Biers, Sarah-Jane Harknett, Stefania Sasso, Simone A. Biagini, Göran Runfeldt, Corinne Duhig, Christopher Evans, Tamsin C. O’Connell, Mait Metspalu, Martin J. Millett, John E. Robb, Toomas Kivisild

## Abstract

The Roman period saw the empire expand across Europe and the Mediterranean, including much of what is today the United Kingdom. While there is written evidence of high mobility into and out of Britain for administrators, traders and the military, the impact of imperialism on local population structure is invisible in the textual record. The extent of genetic change that occurred in Britain before the Early Medieval Period and how closely linked by genetic kinship the local populations were, remains underexplored. Here, using genome-wide data from 52 ancient individuals from Cambridgeshire, we show low levels of genetic ancestry differentiation between Romano-British sites and lower levels of runs of homozygosity over 4 centimorgans (cM than in the Bronze Age and Neolithic. We find fourteen cases of genetic relatedness within and one between sites without evidence of patrilineal dominance and one case of temporary mobility within a family unit during the Late Romano-British period. We also show that the modern patterns of genetic ancestry composition in Modern Britain emerged after the Roman period.

## Introduction

The most visible individuals from Roman Britain through texts are soldiers and administrators, many of whom came from other parts of the Empire (Eckardt and Müldner 2014). The Roman army was recruited from across the whole empire, and, in general, soldiers were posted to areas away from their homelands to avoid conflicts of loyalty (Haynes 2013)(pgs. 121–29). Migration from the Empire into Britain was likely dominated by these groups, as well as traders. Although local populations are archaeologically very well documented, the movement is less well understood and is invisible in the textual record. The extent of mobility in this period has been the subject of recent debate, with work largely focusing on the use of isotope data (Eckardt and Müldner 2014). However, as sampling has been dominated by the examination of unusual burials, our knowledge of the scale of migration and its impact on the overall population is impossible to assess. While the subsequent Early Medieval Period (5th - 10th centuries CE) arguably resulted in major genetic shift towards higher affinities to Dutch, Danish, and other continental North Sea zone ancestries in eastern England, at the scale of 38–75% on average (Schiffels et al. 2016; Gretzinger et al. 2022), it is not clear whether this is due to migration only during the Early Medieval Period or if any of this change could be ascribed to gene flow during the Roman Period and before (Oosthuizen 2017); however, the long-standing ties between Britain and Gaul (Champion 2016)(pg. 155), both prior to and during the Roman period, may obscure the genetic distinction between local, indigenous Britons and incoming individuals.

In contrast to recent genomic studies on demographic changes in Bronze and Iron Age (Patterson et al. 2021) and Early Medieval (Gretzinger et al. 2022) periods, to date, few genomes from Great Britain from the Roman period have been published. A study of seven individuals from a cemetery in York with decapitations showed most individuals had a higher affinity to the modern Welsh than modern English, yet also highlighted the cosmopolitan nature of the Roman empire by identifying an individual with Middle Eastern / North African ancestry (Martiniano et al. 2016). However, York was a cosmopolitan urban centre and cannot be taken as typical of the province as a whole (Ottaway 2004). The population of Roman Britain totalled 2–4 million and was dominated by rural communities, who accounted for about 90% of the people (Millett 1990)(pg. 185). Although extensively networked, these communities were arguably little affected by migration (A. Smith, M. Allen, T. Brindle, M. Fulford 2016)(pg 416–17). However, this hypothesis has not yet been tested. The area that is today Cambridgeshire provides an extensively researched rural, agricultural region, not atypical of the province as a whole, so genetic information from farmstead communities in this region provides a key opportunity to improve our understanding of the local population(s) of Roman Britain.

## Results

To explore the question of migration outside of cosmopolitan centres, we generated genome-wide data for 96 ancient individuals from eight sites (Table 1) in the Cambridgeshire region (Figure 1A) to an average coverage of up to 3.8× (median 0.037×). Mitochondrial haplogroup could be determined for 71 individuals and a subset of 52 genomes which had autosomal coverage >0.01× were used in autosomal analyses including imputation-based analyses relying on 43 genomes (Data S1A). The individuals come from six sites across the Late Iron Age / Romano-British period and two earlier comparative sites from the Neolithic and Bronze Age (Table 1). The majority of our Romano-British sites were in use between 200–400 CE (Table 1, Data S1B) and encompass farmsteads and a cemetery with a number of burials thought to be decapitations (Wiseman et al. 2021) (Supplementary Note 1). In general, the genomes represent an equal distribution of males and females, as determined genetically, and a range of juveniles and adults of all ages (Data S1A). The average endogenous human DNA content varies by site, but overall is 12.03% and genome-wide coverage is 0.13×. Estimated contamination rates from mitochondrial DNA (mtDNA) range between 0 - 3.78% (median 0.83%) and the average misincorporation of C > T in the first five base pairs (bp) is 8.39% (Data S1A).

**Table 1.**
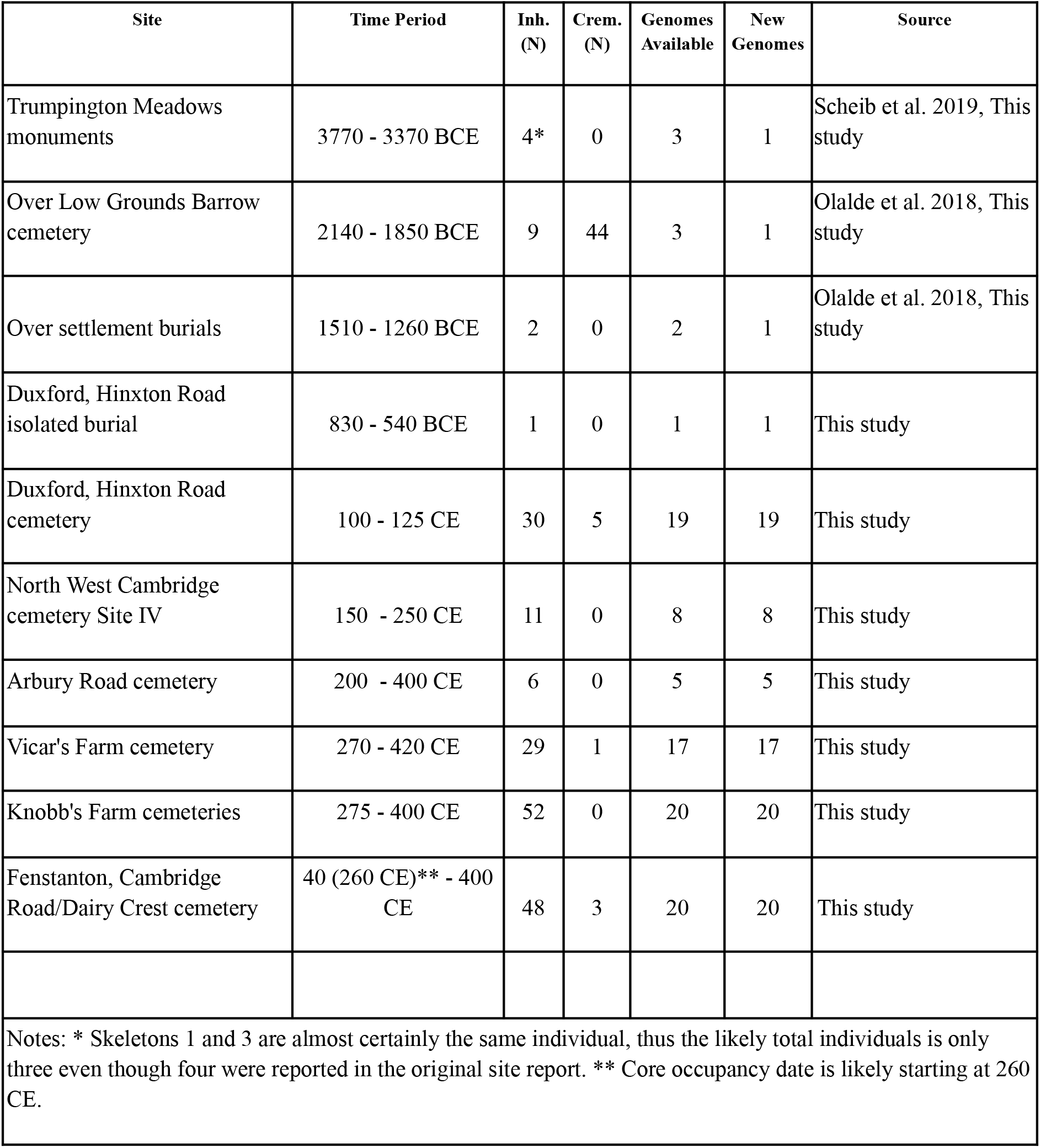
Summary of sites and samples included in this study. Inh. = Inhumations, Crem. = Cremations, Genomes available (N) indicates total individual genomic data including previously published genomes.

**Figure 1.**
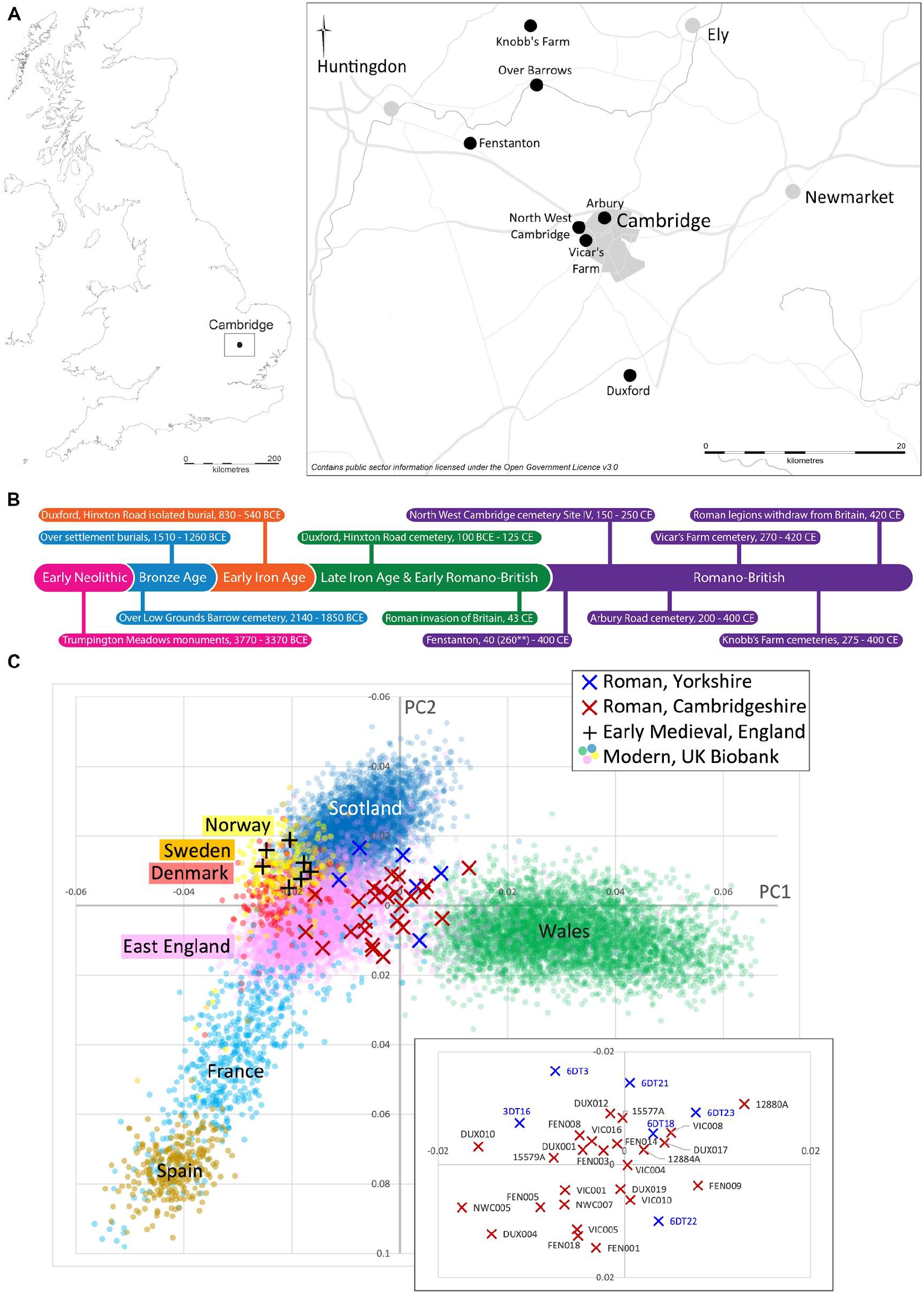
Geographical and chronological distribution of the dataset and population affinities. (A) Site map. (B) timeline. (C) PCA based on imputed genomes.The proportion of total variation explained by PC1 and PC2 is 0.0005 and 0.0004, respectively. Roman and Early Medieval Period genomes shown with x and + symbols, respectively, include those reported in this study as well as those from Martiniano et al. 2016 and Schiffels et al. 2016.

### Population structure of Iron Age / Roman Cambridge

We studied the ancestry of the Roman period genomes from Cambridgeshire in the context of other contemporary material from Britain and modern genomes from Europe and the Middle East using Principal Component Analysis (PCA) (Data S2). We found that Roman genomes from Cambridgeshire all draw their genetic ancestry from Western Europe (Figure 1C) and that similarly to the majority of Roman genomes from Yorkshire they cluster more closely with modern Welsh than local East Anglian genomes (Figure 1C). Unlike the Yorkshire individuals, we do not detect outliers among the 25 Cambridgeshire Roman genomes with >0.1× coverage examined. All Roman period populations examined show homogeneity in their North/West European ancestry in relation to external reference populations in PCA analyses based on imputed data (Figure 1, Figure S1) or projections made from haploid genotype calls (Figures S4 and S5).

We tested whether the imputed Roman genomes have different affinities to ancient and modern European populations using *f*4 statistics. Consistent with the increased Neolithic ancestry observed in Iron Age genomes from England by Patterson et al. (2022), all seven Roman Period sites we tested showed consistently higher drift sharing with Sardinian Neolithic genomes than genomes from Copper and Bronze Age England (-5.3<Z<-2.4; Figure 2A). All sites show higher affinity to Late Iron Age England than to Imperial Roman genomes (Figure 2B). Unlike the Roman Period York cemetery that included a burial of a long distance migrant from present-day Syria or Jordan (Martiniano et al. 2016), we find no evidence of long-distance migration from the Mediterranean region among the 33 imputed genomes from Roman Cambridgeshire that we tested (Figure S2A-C). The Cambridgeshire genomes are also not differentiated by their affinity to Late Iron Age genomes from France, Scotland and England (Figure S2D-E).

**Figure 2.**
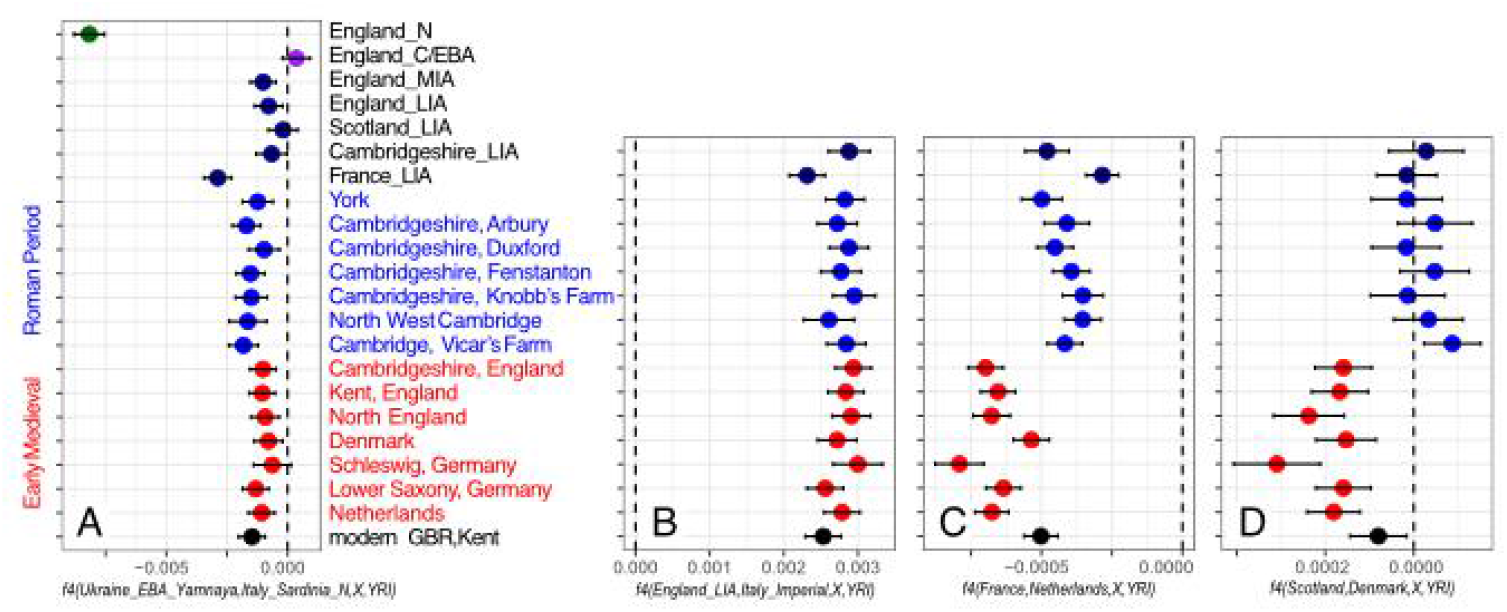
Genetic affinities of Roman Period sites in England to ancient and modern populations of Europe. A-B: affinities to ancient genome groups of individuals from the Allen Ancient DNA Resource v54. C-D: affinities to groups of 200 individuals from the UK Biobank born in France, Netherlands, Denmark and Scotland. Each plot shows the estimated *f4* value with an error range of 2 standard deviations. Respective f4 plots by individuals of the Roman sites are shown in Figures S2-3. Red - Early Medieval Period, blue - Roman Period.

Similarly to Roman Period genomes from York (Martiniano et al. 2016) we find higher proximity of the Cambridgeshire Roman genomes to present-day Dutch than French genomes (Figure 2C, Data S3) and no higher affinity with Danish than Scottish genomes akin to the Early Medieval genomes (Figure 2D). Neither do we observe any notable individual deviations from the patterns observed at site level (Figure S3A-B). In sharp contrast to the single outlier, the long-distance migrant in York, we observe relatively little difference in the affinities of the Roman genomes to present-day groups from England, Kent and East England, with a third of the Roman Period individuals from Cambridgeshire (East England) showing, however, minor but significantly higher affinity to present-day Kent than average present-day genomes from East England (Figure S3C).

We further examined IBD sharing patterns between imputed genomes of Roman individuals from Cambridgeshire in context of available Roman Period data from York, Late Iron Age France (Fischer et al. 2022) and Early Medieval West Europe (Gretzinger et al. 2022) as well as UK Biobank data for individuals born in the UK and elsewhere in Europe (Data S4, Figure 3). Unsurprisingly we find a relatively high level of IBD sharing among geographically close Roman sites in Cambridgeshire with an average probability of 25% of individuals from one site sharing an IBD segment longer than 4cM with individuals from another site, which is more than twice as high as sharing among present-day individuals from East or Southeast England. However, the mean rate of IBD sharing of Cambridgeshire Roman genomes with geographically more distant Roman site from York is lower (23%, p=0.42 by two-tailed t-test) than the local average in context of lower (16%, p=0.0014) diachronic IBD sharing between Cambridgeshire Roman and Early Medieval sites. Notably, IBD sharing among Early Medieval sites from across England is higher (32%, p=0.002) than sharing among Roman sites in Cambridgeshire alone, remaining high for the English Early Medieval sites across the Channel with Early Medieval sites from Lower Saxony and the Netherlands (28%). Compared to Roman sites, the Early Medieval sites from East England show (p=5×10^-7^) increase in IBD sharing with present-day Scandinavian and Dutch genomes from approximately 10% to 15%, which is consistent with the major increase in that period of continental northern European ancestry detected by Gretzinger et al. (2022) (Gretzinger et al. 2022). At the same time, IBD sharing with Late Iron Age France drops in Cambridgeshire from the mean of 15.5% in the Roman to 10% in the Early Medieval and 8% in present-day East England which is comparable to the level of sharing between modern French and English (6.27%) (Data S4).

**Figure 3.**
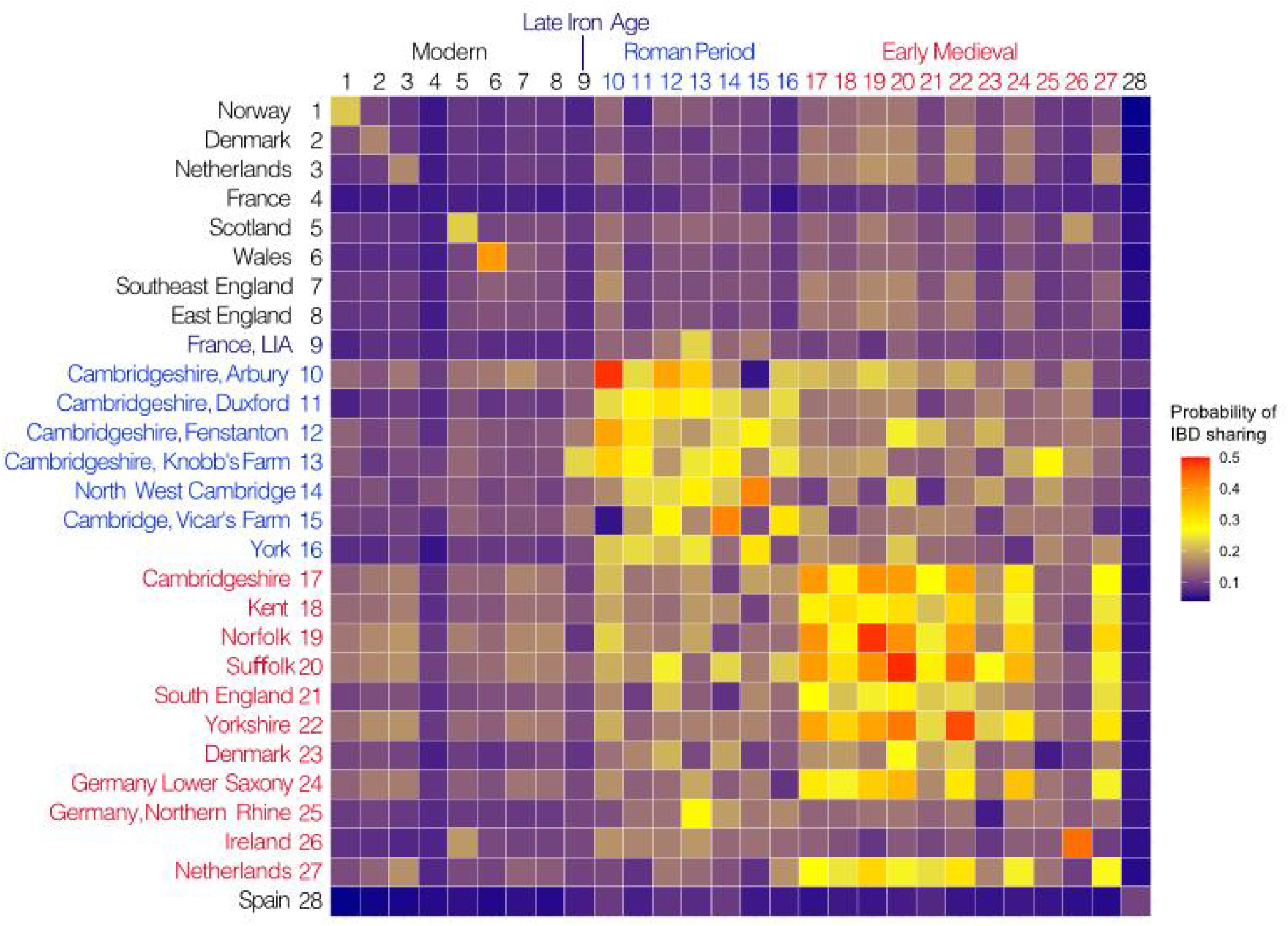
Probabilities of IBD sharing among populations. Heatmap of probabilities of individuals from population i to share at least one IBD segment >4 cM with individuals from population j. Present-day population data from the UK Biobank, ancient imputed genomes include Late Iron Age of France (Fischer et al. 2022), Roman Period data from Cambridgeshire (this study), from York (Martiniano et al. 2016), Early Medieval data (Gretzinger et al. 2022).

We calculated runs of homozygosity (ROH) using HapROH (Ringbauer et al. 2021). Using a two-tailed Student’s T-test we find no difference in the average sum of ROH segments greater than 4cM or 8cM between the Roman era sites (Data S5). Nor, do we find a difference between the two newly generated Bronze Age (Over Barrows) samples and the Roman era populations (Data S5). We do find a difference between our Roman Period populations and the samples from York (4cM p = 0.014; 8cM p = 0.034), but only when comparing our imputed data (haploid mode) to the data available in the Allen Ancient DNA dataset (Anon). When comparing our imputed data to the York ROHs imputed in haploid and diploid mode from the shotgun data, there is no difference detected at either 4cM or 8cM lengths.

### Uniparental marker diversity

To determine variation in the paternal lineages we called the genotypes of 161,140 Y chromosome haplogroup informative binary markers in 29 male samples from Late Iron Age and Roman Cambridgeshire with autosomal coverage >0.01× (Data S1C). All individuals could be assigned to haplogroups common in modern-day Europe. Majority of the samples (85%) belong to haplogroup R1b which became the predominant male lineage in Britain after the spread of the Beaker complex (Data S1D). Two first-degree related individuals from Duxford fall into the I2 clade which captures all previously known Y chromosome lineages in Britain before the Bell Beaker Culture (Data S1D-E). It is not clear, however, whether this particular lineage (I2-Y3722) of the Duxford father-son pair, reflects local continuity and survival from a pre-Beaker population as its present-day distribution is mainly focused on Ireland with only rare cases detected in England and Scotland (https://www.yfull.com/tree/I-Y3722/). Among R1b individuals with more coverage we identify distinct sub-clades, including the British/Irish Bell Beaker signature lineage R1b2-L21 (Patterson et al. 2021) as well as lineages from clades such as R1b11-Z2103 and R1b18-S1194 that have not been encountered in Britain in context of earlier time periods. Notably, none of the four R1b samples with >0.2× Y chromosome coverage fall to the same sub-clade. Some of the identified subclades of R1b appear to be rare in a large, high resolution modern Y chromosome compendium of more than 60,000 FamilyTreeDNA customers (Data S1D). Overall, compared to Copper/Bronze Age periods we do not detect in our Roman Cambridgeshire samples any notable changes in the composition of the Y chromosome haplogroups apart from a single I1 (NWC010) and a single G2a (DUX006) lineage that, by their presence in the Iron Age data by (Patterson et al. 2021), were likely introduced to Britain from the mainland in the Iron Age. (Data S1E). In autosomal DNA and isotope analyses these two individuals (NWC010, DUX006) did not stand out as outliers with external origins or high mobility during their lifetime (Figure 1C, Tables 2-3).

We determined mitochondrial (mtDNA) haplotypes for 71 individuals (Data S1F) and found high diversity, with no haplotype matches except in the case of close kinship (Data S1G).

### Kinship structure

We examined relatedness within and among the Roman sites of Cambridgeshire using a pairwise mismatch approach on pseudo-haploid called data (Monroy Kuhn et al. 2018) to detect 1st-3rd degree related pairs of individuals and an IBD-based approach (Seidman et al. 2020) on imputed genomes to explore more distant forms of relatedness. Despite our relatively small sample sizes per site, we observed closely related pairs (Figure 4) in all Roman age sites from Cambridgeshire except for Knobb’s Farm (Data S1G - H). Both Knobb’s Farm and the previously studied Driffield Terrace (2nd century Roman cemetery in York (Martiniano et al. 2016)), which also did not reveal related pairs, are sites where decapitated burials are common. None of the pairwise comparisons between the sites identified closely related individuals.

**Figure 4.**
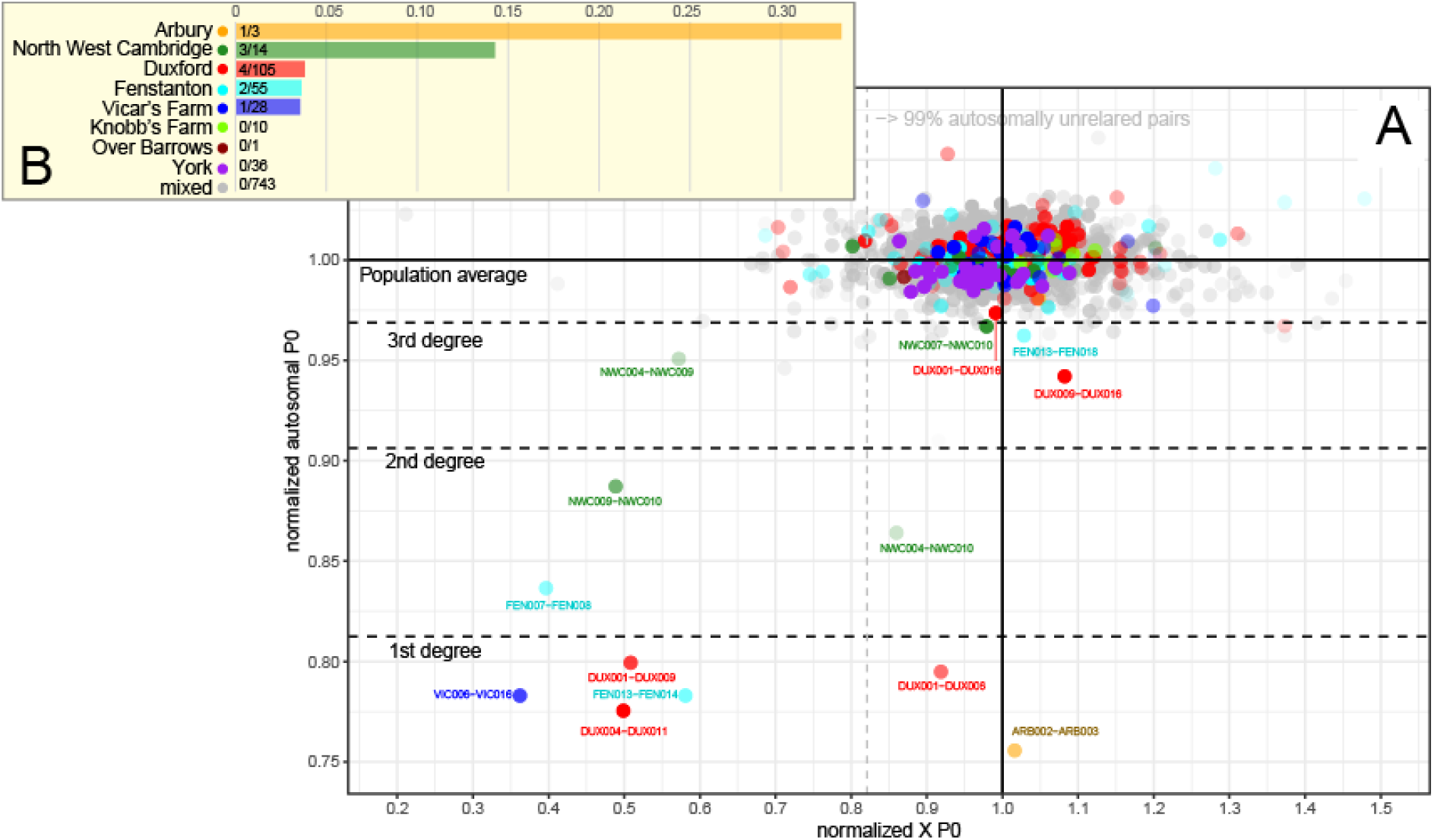
Relatedness between ancient Iron Age/Roman genomes by autosomal and X chromosome calculated in two ways. A) Mismatch probabilities. Each dot shown on the plot represents a pair of ancient genomes assessed (with READ) for their average pairwise differences normalised by population average. First, second and third degree boundaries for the autosomal relatedness are estimated as in Kuhn et al. 2018. The lower boundary for 99% autosomally unrelated pairs is shown on the x-axis for guidance. Dots with high transparency correspond to pairs with low aggregate SNP coverage. B). Proportion of pairs with autosomal 1-3rd degree relatedness per site.

Notably within the relationships detected within the sites we find several triangular cases of relatedness with a female individual involved within more than one pair (e.g. Duxford DUX011 (female) related with DUX019 (male) and DUX001 (male)), or in case of North West Cambridge we find a relationship between three sampled male individuals (NWC004, NWC010, and NWC009) who appear to be related to each other through unsampled female(s) (either unexcavated or not sampled) because their pairwise X chromosomal differences are lower than population average despite the fact that they carry different mtDNA lineages (Figure 4). Genetically related individuals appear not to be clustered or buried next to each other: for example, the members of a Duxford family DUX011 (mother), DUX008 (father) and their son (DUX001) are all buried in different groups of burials (Figure S6) identified in the original site report (Lyons 2011). Or, similarly, in Vicar’s Farm related pairs of individuals were buried in different groups of burials (Evans and Lucas 2020) (pages 333-34 & 377).

We further used IBIS to explore IBD sharing within and among Late Iron Age and Roman sites in Cambridgeshire. In case of all pairs of imputed individuals that were identified with READ as closely related we found multiple IBD segments supporting their close relatedness (Data S4). However, in all cases the observed total IBD shared expressed in kinship coefficient was less than expected from the 1st-3rd degree relationship suggesting that capturing long tracts of IBD at low coverage is hindered by their fragmentation due to imputation errors. Besides the kinship pairs already detected with READ (Figure 4) we did not find any new relationships with IBIS within the sites. However, we detected a case of distant relatedness between DUX019 from Duxford and a previously reported sample 12884A (HI2, Schiffels et al. 2016) from Hinxton, who share five IBD segments longer than 7cM consistent with estimated kinship coefficient suggesting 6th degree relatedness. Given the Duxford and Hinxton sites are located only 3 kilometres from each other and are both in the Cam valley this finding points to local mobility between geographically adjacent sites (Data S4).

### Phenotypic Changes / Health

To investigate potential changes in Cambridgeshire through the Roman Period, we imputed 114 SNPs known to be involved in phenotypic traits related to diet (carbohydrate, lipid and vitamin metabolism), immunity (response to pathogens, autoimmune diseases and other immunity traits) and pigmentation (eye, hair and skin colour) in the ancient individuals presented here plus 234 individuals from the literature, for a total of 277 samples divided into four groups from the Mesolithic to the Roman Period (Data S6A-D). Within Great Britain, from the Neolithic to modern Great Britain (1000 Genomes GBR) there are 34 significant SNPs (Data S6B). We can see two main “break-points” considering the significant group pairs after the Tukey test for each of the 34 SNPs, in line with previous findings: 1) after Neolithic and 2) after the Bronze Age. Most of the significant SNPs involve Neolithic or Chalcolithic/Bronze Age groups that differ from later periods. More specifically, they are involved in several different diet, immunity and pigmentation functions. In the diet group, we found six significant SNPs: two that confer lactase persistence, one involved in lipid metabolism, two in fatty acid metabolism and one in vitamin D metabolism. The SNP involved in the lipid metabolism is in an introgressed Denisovan-tract and tends to increase over time, with greater shifts after Neolithic and Chalcolithic/Bronze Age. The fatty acid metabolism SNPs show a change in frequency mainly after Chalcolithic/Bronze Age, while the SNP that is a protective factor against vitamin D deficiency show the lowest frequency during Chalcolithic/Bronze Age (0.49 vs. 0.67 in Neolithic and 0.72 on average in previous and later periods respectively).

When focusing on the group pairs involving Iron Age/Romans (IAR), we see eight SNPs are different between IAR and modern GBR. The MCM6 locus, with two Lactase-persistence SNPs, show a sharp increase after Iron Age/Romans, after the first great increase after Bronze Age; this is consistent with recent findings related to low frequency of LCT alleles in Bronze Age and increase in frequency in later periods (e.g. (Burger et al. 2020; Segurel et al. 2020)). Between different sites of Roman UK and Roman Italy, no significant SNPs appear. Differently from (Kerner et al. 2021), we do not observe frequency fluctuations for the TB risk factor rs34536443, which is low in frequency from the Neolithic with no significant changes over time. It reached present-day frequency from IAR.

### Mobility through isotopic analysis

To further explore potential childhood origins and geographical mobility, we measured oxygen isotope ratios in the tooth enamel of Iron Age and Roman individuals excavated from the Cambridgeshire region. The oxygen isotope composition of local water sources is largely determined by the local climatic conditions (Dansgaard 1964; Pederzani and Britton 2019). The oxygen isotope ratios measured in archaeological human tooth enamel are a reflection of the water consumed during the formation of the enamel during childhood and can therefore provide information about the environment the individual grew up in (DeNiro and Epstein 1978; Longinelli 1984; Luz, Kolodny, and Horowitz 1984; Luz, Kolodny, and Kovach 1984). This can be broadly used to identify population mobility if this does not match the expected values for the environment in which they are buried (Pederzani and Britton 2019).

We measured the carbonate oxygen isotope ratios (δ^18^O_CO3_) of 33 second premolars from 17 individuals (1 Early Iron Age, 1 Middle Iron Age, 15 Late Iron Age/Early Roman) from Duxford and 15 individuals (all Mid-Late Roman) from Vicar’s Farm, Cambridge. We compared the results to published δ^18^O_CO3_ data from 33 individuals from Knobb’s Farm (1 Middle Iron Age, 32 Late Roman: (Wiseman et al. 2021). The datasets are broadly comparable, although due to the variety of teeth analysed, the data will not represent exactly the same period of life (Lightfoot in Wiseman et al. 2021, p. 160).

The δ^18^O_CO3_ values across the three sites are wide-ranging and overlapping (Figure 5) (Data S7). The distribution of the δ^18^O data slightly differs. However, the range of values seen at Duxford and Vicar’s Farm fit well within the distribution of the data from Knobb’s Farm (Figure 5).

**Figure 5:**
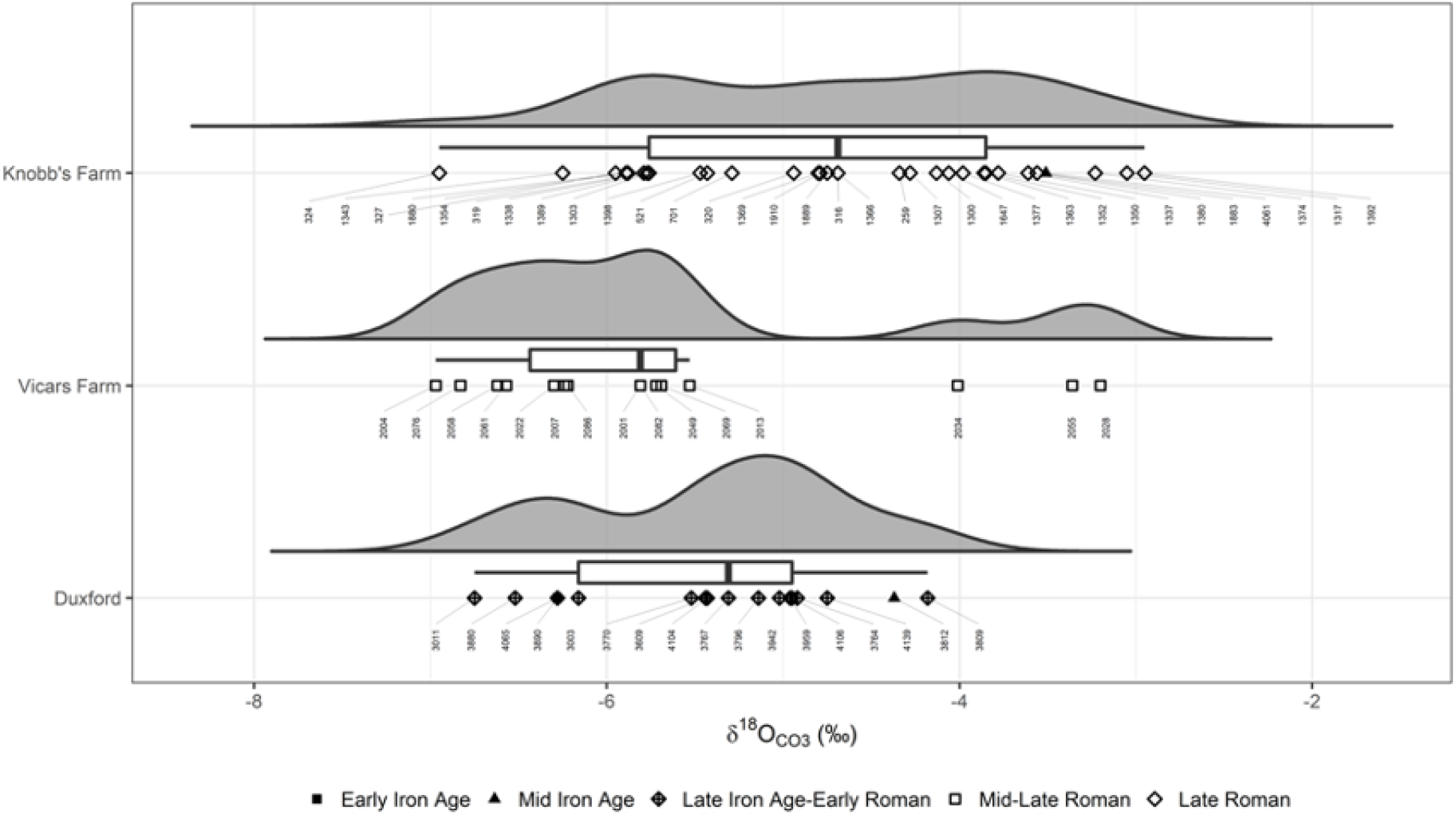
**Raincloud plot of δ18OCO3 values from Duxford, Vicars Farm and Knobbs Farm**, showing probability distribution, median, IQR, outliers and scatter of data, with individual skeleton numbers. Data for Knobb’s Farm sourced from Wiseman et al. (2021).

Converting the δ^18^O_CO3_ values to phosphate oxygen isotope values (δ^18^O_PO4_) (Coplen 1988; Chenery et al. 2012) allows for broad comparisons with previously published data and expected ‘local’ environmental values (Figure 5). The mean expected ‘local’ range of δ^18^O_PO4_ values for the eastern low rainfall zone, in which the three sites are located, has been estimated at 17.2‰ ±1.3 (2SD) (Evans et al. 2012). The isotope values for Knobb’s Farm (mean: 17.2‰ ±2.2) fits well with this estimate, while the values for Duxford (mean: 16.5‰ ±1.6) and Vicars Farm (mean: 16.2‰ ±2.6) are slightly lower but are still within the general range. Only skeleton 2004 (δ^18^O_PO4_ = 14.8‰), a Mid-Late Roman male from Vicar’s Farm and skeletons 324 (δ^18^O_PO4_ = 14.8‰), a Late Roman male (genetics = XY, macroscopic sex=?female) and 1392 (δ^18^O_PO4_ = 19.1‰), another Late Roman male, from Knobb’s Farm have δ^18^O_PO4_ values which are on the edge or beyond the overall total range of ‘local’ values currently estimated for Britain (Evans et al. 2012, Lightfoot and O’Connell 2016).

We investigated the presence of potential outliers further, following Lightfoot and O’Connell (2016) (Table 3)). The 1.5IQR method is considered most robust in this instance and identifies outliers only at Vicar’s Farm: skeletons 2028, 2034 and 2055. However, these are well within the overall range of values seen at Knobb’s Farm and may only appear as outliers due to small sample sizes. Interestingly, the individuals identified as outliers using 1.5IQR are not the same as those highlighted above as being outside the expected δ^18^O_PO4_ range, highlighting the difficulties of robustly identifying and interpreting potential childhood origins and geographical mobility using oxygen isotopes alone.

Statistical comparisons of all sampled individuals from the three sites indicate that the samples were unlikely to be taken from populations with the same distributions (Kruskal-Wallis δ^18^O_CO3_: p=0.005, es=0.139), with the differences lying between Vicars Farm and Knobb’s Farm (Dunn’s post-hoc with Bonferroni adj: p-adj=0.007). For individuals that were confidently assigned a sex estimate of female or male, when both sex and site are considered, sex does not appear to correlate with δ^18^O_CO3_ values, but site does (Two-way ANOVA δ^18^O_CO3_ (site): p=0.033, es=0.119; δ^18^O_CO3_: p=0.803, es=0.001). There also appears to be no difference in the populations by time period when the individuals were assigned to broad date categories of Iron Age (incorporating those dated Early and Mid-Iron Age), Late Iron Age-Early Roman, and Roman (incorporating those dated to Mid-Late and Late Roman), (Kruskal-Wallis δ^18^O_CO3_: p=0.516, es=-0.010).

## Discussion

The region of Cambridgeshire appears to have been composed mostly of homogenous, local populations when compared to the York samples of the same period where one out of seven randomly sampled individuals was a long-distance migrant. This evidence supports the hypothesis that the rural populations of Roman Britain were largely unaffected by migration. The differences between PCA mapping of Roman-period populations closer to Celtic populations and PiC score-based analyses show more similar regional affinities across different British geographic regions to Cambridgeshire Roman Period. This may potentially be explained by the relative homogeneity of the allele frequency pool before Anglo-Saxon migrations that makes the allele frequency approach more sensitive to detect the differences.

Regardless of relatively small sample sizes, we find close pairs of relatives in most of the Cambridgeshire sites. The exception being Knobb’s Farm, a cemetery associated with a settlement that was possibly engaged in the processing of agricultural products and in which there were a significant number of burials that were decapitations. Knobb’s Farm appears to have been more broadly networked than the other local farming communities sampled here, yet distinct from the cosmopolitan urban centre at York, which may explain the difference in population heterogeneity.

Generally, the oxygen isotope range of individuals from the three sites is wide, but falls within the range of previous archaeological data from the UK (Lightfoot and O’Connell 2016) and from Cambridgeshire specifically (Rose 2020). The δ^18^O_PO4_ values from all three sites are broadly within those expected for the British eastern bioclimatic zone (Evans et al. 2012), indicating that the majority of the individuals studied could have spent their childhoods in the local area, or at least, an area with similar climatic conditions to Cambridgeshire. Six individuals out of 33 can be considered different to the others. Three individuals are different to the rest in terms of the expected values for this region: skeleton 2004 from Vicar’s Farm and skeletons 324 and 1392 from Knobb’s Farm exhibit δ^18^O_PO4_ values which fall above or below expected ‘local’ values (Evans et al. 2012; Lightfoot and O’Connell 2016; Wiseman et al. 2021) which could indicate that they spent their childhoods outside Cambridgeshire (Table 3). These may be the most likely candidates for being ‘non-locals’ – skeletons 2004 and 324 may have spent their childhoods somewhere with a colder climate than Cambridgeshire and skeleton 1392 may have spent their childhood in a warmer, drier environment (Wiseman et al. 2021).

Three others, skeletons 2028, 2034 and 2055 from Vicar’s Farm, are statistical outliers for δ^18^O_CO3_ compared to the rest. This may indicate that these three individuals spent time in a geographically/climatically distinct area to the rest of the group when their teeth were developing. This could be particularly interesting as aDNA analysis identified skeletons 2028 and 2076 as likely to be brothers, however, their δ^18^O_CO3_ values are very different (skeleton 2028 = -3.20‰, skeleton 2076= -6.83‰), with skeleton 2028 having the highest δ^18^O_CO3_ value at Vicar’s Farm. This could indicate that the brothers were not raised in the same geographical location. However, the overall range of the Vicar’s Farm δ^18^O_CO3_ values is very similar to the other two sites and it is quite possible that the apparent bimodality is a by-product of the small sample size, and that if a larger number of samples had been analysed from the site, the distribution would be more normal and the difference between the brothers could be considered part of ‘normal variation’ at the site.

Overall, the results of the oxygen isotope analysis of the individuals from Duxford, Vicar’s Farm and Knobb’s Farm are fairly homogeneous and indicate that the majority of the population were likely to have spent their childhoods either in the local area, or in an area with similar climatic conditions to Cambridgeshire. There is no correlation between the oxygen isotope results and key demographic or burial information. There is no difference in oxygen isotope values between females and males at each site nor by time period, meaning there is no isotopic evidence for a large-scale change in population diversity (i.e. differences in the number of individuals who may have originated from elsewhere) between the Iron Age and Roman period sites. The data does appear to highlight several individuals that may have spent their childhoods elsewhere, however, the difficulties of interpreting δ^18^O data due to large ranges and broad, overlapping estimations of expected values means that these individuals can only be tentatively identified as potential migrants and further corroborating evidence, such as strontium isotope analysis would be required to make a more definitive interpretation.

## Materials and Methods

### Sample information and ethical statement

All skeletal elements were sampled with permissions from the representative bodies/host institutions. Samples were taken and processed to maximise research value and minimise destructive sampling. Teeth were sampled from skeletons using gloves. Molars were preferred due to having more roots and larger mass, but premolars were also sampled.

### Archaeological sites and material

#### Over Barrows

At the Over Low Grounds site 13 km northwest of Cambridge, excavated in 2008 by the Cambridge Archaeological Unit, a small Beaker period cemetery of six inhumations underlay a collared urn-associated Early Bronze Age barrow cemetery. Two of these individuals were dated to 2199–1960 and 2126–1912 cal BC and the earliest is likely to have been buried 2140–1970 cal BC (Evans, C., Tabor, J., and Vander Linden, M., 2016, pp. 336–7). There were also some later burials of neonates dated c. 1900–1850 cal BC. Nearby were two Middle Bronze Age inhumation burials within a settlement, dated to 1511–1303 and 1449–1260 cal BC (Evans, C., Tabor, J., and Vander Linden, M., 2016, p. 253) (Evans et al. 2016).

#### Duxford

The site off Hinxton Road, Duxford, was excavated by CAM ARC in 2002 (Lyons 2011). Located 11km southeast of Cambridge, it is situated on a chalk knoll overlooking a crossing of the River Granta, a tributary of the River Cam. There was an Early Iron Age crouched inhumation dated to 827–540 cal BC and two supposedly Middle Iron Age inhumations, one dated to 386–111 cal BC (Lyons, A., 2011, pp. 10–12, 15–16), although aDNA analysis presented here indicates that these may be Late Iron Age. During the Late Iron Age, the higher ground was defined by a series of ditches that were repeatedly redug, surrounding a short-lived timber-framed rectangular shrine and a burial ground that was in operation c. 100 CE – 125 CE (Lyons, A., 2011, pp. 38–49). The burials are believed to have ‘formed a selected part of a community perhaps largely made up of a single family or other social grouping’ (Lyons, A., 2011, p. 38). A range of orientations and grave goods were present, with the 27 or more burials containing 37–8 individuals divided into four or five groups based on spatial patterning, orientation etc. (Group 1a six inhumations and three cremations; Group 1b two cremations; Group 2 nine inhumations; Group 3 three inhumations; Group 4: six inhumations).

#### Vicar’s Farm

Vicar’s Farm is a rural settlement located 1.3km west of the extensive Romano-British roadside settlement of Cambridge, falling in its immediate hinterland. Excavated by the Cambridge Archaeological Unit in 1999–2000 (Evans and Lucas 2020), there is evidence for Iron Age activity with a Romano-British settlement that commences c. 80 AD with a cremation cemetery, a small timber shrine and a farmstead with a rectilinear ditch system, aisled building and various other enclosures. Over time the settlement expanded and c. 270 AD an inhumation cemetery was established on the southern edge of the settlement within the ditched enclosure system (Evans, C. and Lucas, G., 2020, pp. 314–37) (Fig. 3.46). This cemetery presumably served part or all of the nearby rural settlement, which shows some signs of being of higher status than most other local settlements and may have fulfilled some minor central place role within the local rural settlement hierarchy. There is evidence that neonates were buried within the settlement itself rather than the cemetery and it is possible that high status individuals were buried elsewhere.

The studied skeletons come from the inhumation cemetery, where thirty individuals were recovered from 29 graves. Eight individuals appear to have been buried in coffins, while hobnails indicate that seven were either wearing or accompanied by footwear. Grave goods accompanying seven individuals included bracelets, finger-rings, a glass bead necklace, ceramic vessels and a cache of glass fragments. Most graves are orientated roughly north–south or east–west and the burials were mainly extended and supine, with just one crouched burial.

#### North West Cambridge

Archaeological investigations at North West Cambridge by the Cambridge Archaeological Unit between 2009 and 2019 revealed a series of rural Romano-British settlements. The sampled skeletons come from settlement RB2.C (Site IV) (Cessford and Evans 2014). Initially crossed by a double-ditched boundary the area was initially largely empty until it was divided into a series of ditched enclosures. An inhumation cemetery was established within one of the enclosures. This consisted of eleven definite and one possible burials, plus another burial a short distance away. These were largely of adults with some possible sub-adults and span the period c. 150–250 CE, although burials may have continued slightly after that time. Eleven of the burials had some evidence for coffins, ten or eleven of the burials had hobnailed shoes and five or six were accompanied by beakers. There may have been some other grave goods although these are less certain, and there was a single decapitation burial.

#### Knobb’s Farm

Excavations at Knobb’s Farm, Somersham, Cambridgeshire, by the Cambridge Archaeological Unit between 2000 and 2010 uncovered three small late Roman cemeteries, positioned at the edge of a farming settlement by boundary ditches in a former field system dating to the fourth century CE (Wiseman et al. 2021). The 52 burials found (11 individuals from eight graves in Cemetery 1; 28 individuals from 30 graves in Cemetery 2; 13 individuals from 12 graves in Cemetery 3) included 17 decapitated bodies and 13 prone burials. At least three bodies were buried in coffins, 15 were accompanied by pottery vessels with other grave goods including an antler comb, 30 beads and the remains of a box. It has been suggested that the decapitated burials relate to judicial execution.

#### Fenstanton Cambridge Road (CRF2345) and Fenstanton Dairy Crest (DC3112)

Albion Archaeology evaluated and then dug two adjacent sites at the southern edge of the village of Fenstanton, 15 km north-west of Cambridge and close to the Via Devana. The River Great Ouse runs 1.5km to the north of the village, and the underlying geology is sand and gravels overlying mudstone. The Cambridge Road site is primarily on level pasture and lies at a height of c. 7m OD; Dairy Crest is on a former dairy site with modern buildings and hardstanding, c. 4–15 m OD.

The open-area excavations of 2017–2018 (c. 5.5 ha excavated) revealed the area had late Iron Age material succeeded by a large enclosed settlement, occupied from the beginning of the Roman period and continuing into the latter half of the 4th century; there were traces of a Late Roman timber building. It was probably primarily agricultural and contained a specialist cattle butchery and evidence of domestic, craft and small-scale industrial activity; some above-average status occupation is suggested by fine ware, high-status artefacts and building ceramics.

Several clusters of inhumations were found, in total containing 48 individuals, plus three cremations including a bustum. Graves were primarily NW-SE, inhumations extended or semi-flexed supine but some non-normative (prone, contracted, splayed knees, head to SE). Many nails were within graves, suggesting coffins or biers, together with dress accessories and hobnail. One burial has apparent evidence for crucifixion.

#### Arbury

The burials were discovered in August of 1952 during construction work (Fell 1956) on Fortescue Rd, Arbury. One lead-lined coffin containing a female skeleton inspired the poem, “All the Dead Dears” by Sylvia Plath. Up to six burials were uncovered (Fell 1956).

### Generation and analysis of Isotopic data

For this study, teeth from 17 individuals (1 Early Iron Age, 1 Middle Iron Age, 15 Late Iron Age/Early Roman) from Hinxton Road, Duxford and 15 individuals (all Mid-Late Roman) from Vicar’s Farm, Cambridge were sampled for carbonate δ^18^O analysis (δ^18^O_CO3_). Only permanent second premolars (PM2) or second molars (M2) were selected for analysis, with the enamel development of these teeth occurring between c.2.5-7.5yr (AlQahtani 2008).

Pre-treatment of the enamel samples was carried out following a protocol based on methods in Balasse et al. (2002) (Balasse et al. 2002). To remove surface contaminants, the outer surface of the tooth enamel was abraded using a handheld Dremel drill with a round-headed, diamond-tipped drill bit. Following this, approximately 5.5-10.0mg of enamel powder was collected using a smaller round-headed diamond-tipped drill bit. Samples were then vortex mixed in approximately 0.1ml per mg of sample of 2-3% aq. sodium hypochlorite (NaOCl) and refrigerated for 24h. Samples were rinsed five times with distilled water and then vortex mixed in 0.1ml per mg of sample of 0.1M aq. acetic acid (CH_3_COOH) and left at room temperature for 4 h. Samples were then rinsed five times with distilled water, frozen and placed in a freeze dryer until full lyophilization. Approximately 2-4mg of the resultant enamel powder was weighed into glass gas bench tubes. For each batch of samples submitted for analysis, 2-4mg of two in-house faunal enamel standards were also weighed into glass gas bench tubes (eight standard tubes in total). The glass vials were vacuum sealed and the samples were reacted with 100% orthophosphoric acid at 90°C using a Micromass Multicarb Sample Preparation System. The CO_2_ produced was then dried and transferred cryogenically into a Gas Bench II coupled to a Delta V mass spectrometer in the Godwin Laboratory, Department of Earth Sciences, Cambridge. All results for both carbon and oxygen are measured and reported on the international scale relative to VPDB calibrated through the NBS19 standard (Coplen 1988; Hoefs). Based on repeated measurements of the international and in-house standards, analytical error was < ± 0.10‰ for δ^18^O_CO3_.

All 1^18^O values are primarily reported as δ^18^O_CO3_ (VPDB) values. Phosphate 1^18^O (1^18^O_PO4_) (VPDB) values have been estimated by converting δ^18^O_CO3_ (VPDB) to δ^18^O_CO3_ (VSMOW) using equation: δ^18^O_CO3_ (VSMOW) = 1.03091 × δ^18^O_CO3_ (VPDB) + 30.91 (Coplen 1988). Then converting δ^18^O_CO3_ (VSMOW) to δ^18^O_PO4_ (VSMOW) using equation: δ^18^O_PO4_ (VSMOW) = 1.0322 × δ^18^O_CO3_ (VSMOW) – 9.6849 (Chenery et al. 2012). δ^18^O_PO4_ (VSMOW) values are more comparable with other datasets but each conversion does incur error (Chenery et al., 2012).

All statistical analysis and graphical representations of the results were performed using R version 4.0.3 and R Studio version 1.4.1106. Statistical analysis was primarily undertaken using R package ‘rstatix’, following Kassambara (2019) (Kassambara 2019). Where p-value based null hypothesis testing was used, appropriate testing of assumptions was carried out to make sure there were no major violations of the methods and non-parametric testing was applied where appropriate. Any p-values generated were considered in context and making conclusions drawn primarily from p-values alone was avoided. Outliers were identified using three methods: >1.5 × IQR, >3 × median absolute deviations (MAD) from median and >2 x standard deviation (SD) from mean (see (Lightfoot and O’Connell 2016)). Raincloud plots were produced following Allen et al. (2019) (Allen et al. 2019), using R code by Allen et al., and the R package ‘cowplot’. Raincloud plots combine a ‘split-half violin’ plot (showing the probability density), a boxplot (showing the median and interquartile range (IQR)) and a jittered raw data scatterplot.

### Sampling, ancient DNA extraction and library preparation

Samples were processed in the clean room of the dedicated ancient DNA laboratory of the Institute of Genomics, University of Tartu, Estonia following established protocols as detailed most recently in (Saupe et al. 2021).

### DNA sequencing

DNA was sequenced using the Illumina NextSeq500/550 High-Output single-end 75 cycle kit. As a norm, 15-20 samples were sequenced together on one flow cell; additional data was generated for 34 samples to increase coverage (Data S1A).

### Mapping

Before mapping, the sequences of the adapters, indexes, and poly-G tales occuring due to the specifics of the NextSeq 500 technology were cut from the ends of DNA sequences using cutadapt-1.11 (Martin 2011). Sequences shorter than 30 bp were also removed with the same program to avoid random mapping of sequences from other species.

The sequences were aligned to the reference sequence GRCh37 (hg19) using Burrows-Wheeler Aligner (BWA 0.7.12) (Li and Durbin 2010) and the command *aln* with re-seeding disabled.

After alignment, the sequences were converted to BAM format and only sequences that mapped to the human genome were kept with samtools 1.3 (Li et al. 2009). Afterwards, the data from different flow cell lanes were merged and duplicates were removed using picard 2.12 (http://broadinstitute.github.io/picard/index.html).

### aDNA authentication

As a result of degradation over time, aDNA can be distinguished from modern DNA by certain characteristics: short fragments and a high frequency of C=>T substitutions at the 5’ ends of sequences due to cytosine deamination. The program mapDamage2.0 (Jónsson et al. 2013) was used to estimate the frequency of 5’ C=>T transitions. Rates of contamination were estimated on mitochondrial DNA by calculating the percentage of non-consensus bases at haplogroup-defining positions as detailed in (Jones et al. 2017). Each sample was mapped against the RSRS downloaded from phylotree.org and checked against haplogroup-defining sites for the sample-specific haplogroup.

Samtools 1.3 (Li et al. 2009) option *stats* was used to determine the number of final reads, average read length, average coverage etc. The average endogenous DNA content (proportion of reads mapping to the human genome) was 12.03% (0.003 - 54.65%).

### Calculating genetic sex estimation

Genetic sex was calculated using the methods and script described in (Skoglund et al. 2013), estimating the fraction of reads mapping to Y chromosome out of all reads mapping to either X or Y chromosome. Genetic sex was calculated for samples with a coverage >0.01× and only reads with a mapping quality >30 were counted for the autosomal, X, and Y chromosome.

### Determining mtDNA haplogroups

Mitochondrial DNA haplogroups were determined using Haplogrep2 on the command line (Kloss-Brandstätter et al. 2011). Subsequently, the identical results between the individuals were checked visually by aligning mapped reads to the reference sequence using samtools-1.3 (Li et al. 2009) command tview and confirming the haplogroup assignment in PhyloTree (accessed at: www.phylotree.org). Additionally, private mutations were noted for further kinship analysis.

### Y chromosome variant calling and haplotyping

A total of 161,140 binary Y chromosome SNPs that have been detected as polymorphic in previous high coverage whole Y chromosome sequencing studies (Hallast et al. 2015; Karmin et al. 2015; Poznik et al. 2016) were called in 29 male samples with more than 0.01× autosomal coverage using ANGSD-0.916 (Korneliussen et al. 2014) ‘-doHaploCall’ option. A subset of 144,550 sites yielded a call in at least one of the samples and in the case of 5653 sites at least one of the 29 samples carried a derived allele (Table S8). Basal haplogroup affiliations (Table S7) of each sample were determined by assessing the proportion of derived allele calls (pD) in a set of primary (A, B, C…T) haplogroup defining internal branches, as defined in (Karmin et al. 2015), using 1677 informative sites. In case of 25/29 samples (with the exception of the four lowest coverage samples whose haplogroup affiliation could only be supported by two sites) the primary haplogroup could be determined unambiguously with the support of at least 3 variants in the derived state. Further detailed sub-haplogroup assignments within the phylogeny of the primary haplogroup were determined on the basis of mapping the derived allele calls to the internal branches of the FamilyTreeDNA tree based on approximately 52,500 modern high coverage genomes (sequenced with the Big Y technology) and highlighting the marker tagging the branch with the lowest derived allele frequency (Table S7).

### Preparing the datasets for autosomal analysis

Autosomal variants were called with the ANGSD-0.921 software (Korneliussen et al. 2014) command --doHaploCall keeping base for the 1,233,534 positions that are present in the 1240K capture set. Two compiled datasets were downloaded from David Reich Lab (https://reich.hms.harvard.edu/downloadable-genotypes-present-day-and-ancient-dna-data-compiled-published-papers, release: March 1 2020): 1) “1240K” consisting of 3589 ancient, 2933 present-day individuals covered at 1,233,534 positions and 2) “1240K + HO” consisting of 3589 ancient, 6472 present-day individuals covered at 597,573 positions.

Samples from this study were merged with the comparative dataset using PLINK v1.90 (Chang et al. 2015) and in the resulting merged file #1, SNPS were filtered for a genotyping rate over 10% (--geno 0.9) leaving 1,232,632 SNPs from the base dataset. In file #2, SNPS were filtered for a genotyping rate over 10% (--geno 0.9) leaving 597,573 SNPs from the base dataset. Both filesets were converted to EIGENSTRAT format using the program *convertf* from the EIGENSOFT 7.2.0 package (Patterson et al. 2006).

### Principal component analysis

PCA was performed using the program smartpca from the same package, projecting ancient individuals onto the components constructed based on the modern genotypes using autoshrink:YES and lsqproject: YES.

### Kinship analysis

Variants were called with ANGSD (Korneliussen et al. 2014) command --doHaploCall, sampling a random base for the positions that are present at MAF>0.1 in the 1000 Genomes GBR population (1000 Genomes Project Consortium et al. 2015) giving a total of 5,499,776 SNPs for autosomal and 158,249 SNPs with MAF>0.05 in UK10K males for X chromosome analyses. For the comparison with published studies, the merged plink file from the PCA analysis was used and select populations retained using plink --keep and converted to .tped. The ANGSD output files were converted to .tped format, which was used as an input for kinship analyses with READ (Monroy Kuhn et al. 2018). When using this approach the sample size and population diversity is important, too small a sample size will shift the estimated P0 upwards lead to false negatives (not detecting 1st or 2nd degree relatives) and too much population diversity, i.e. analysing completely different populations together (separated by too much time, distance etc.) will shift the P0 values lower, leading to false positives. Keeping this in mind we analysed the data in several ways according to geography and time. First all Roman period sites were analysed together, then each site separately. Identical individuals and first degree relatives are consistently detected across groupings (Data S1H). In addition to 1st and 2nd degree relationships we also estimated P0 cutoffs (15/16 = 0.9375 as per Kuhn et al. 2018) for the detection of 3rd degree relatives while acknowledging that due the lack of relevant empirical data we cannot estimate the error rates for this class of relationship.

Imputed genomes were used to study in further detail cases of close degree of genetic relatedness detected with READ. We used the --genome function of PLINK 1.9.0 (Chang et al. 2015) to estimate pairwise proportions of IBD1 and IBD2 that are informative, for example, for distinguishing parent-offspring from sibling relationships (Data S1G).

### Genome imputation

Following (Hui et al. 2020), genotype likelihoods were first updated with BEAGLE 4.1 (Browning and Browning 2016) from genotype likelihoods produced by ATLAS (Link et al. 2017) in Beagle -gl mode, followed by imputation in Beagle -gt mode with BEAGLE 5 (Browning et al. 2018) from sites where the genotype probability (GP) of the most likely genotype reaches 0.99. To balance between imputation time and accuracy, we used 503 Europeans genomes in 1000 Genomes Project Phase 3 (1000 Genomes Project Consortium et al. 2015) as the reference panel in Beagle -gl step, and 27,165 genomes (except for chromosome 1, where the sample size is reduced to 22,691 due to a processing issue in the release) from the Haplotype Reference Consortium (HRC) (McCarthy et al. 2016) in the Beagle -gt step. A second GP filter (MAX(GP)>=0.99) was applied after imputation. Because Beagle treats “./.” in the VCF input as sporadically missing and imputes them during haplotype phasing, which damages the accuracy when such missing genotypes are common, we imputed each genome individually so that missing genotypes were not included in the VCF input to Beagle 5. These include:

All new genomes sequenced in this study; Mesolithic genomes (n=3), Neolithic and Chalcolithic genomes (n=13), Iron Age and later genomes (n=11) from (Antonio et al. 2019); Bronze Age Steppe: RISE509, RISE511, RISE547, RISE548, RISE550, RISE552 from (Allentoft et al. 2015); EBA1 and EBA2 from (Damgaard et al. 2018); Anatolia Neolithic: Bar31, Bar8, Klei10, Pal7, Rev5 from (Hofmanová et al. 2016); WHG: KO1 from (Gamba et al. 2014); Bichon from (Jones et al. 2015); La Braña from (Olalde et al. 2014); EHG: Sidelkino from (Damgaard et al. 2018); CHG: KK1 and SATP from (Jones et al. 2015).

### Runs of Homozygosity

We used hapROH (Ringbauer et al. 2020) to detect runs of homozygosity (ROH) in ancient genomes. Using information from a reference panel, hapROH has been shown to work for genomes with more than 400K of the 1240K SNPs panel covered at an error rate lower than 3% in pseudo-haploid genotypes (Ringbauer et al. 2020). We note that the requirement is broadly in line with the imputation accuracy we get from coverages as low as 0.05×, where ∼60% of common variants (MAF≥0.05) in the HRC panel are recovered with an accuracy greater than 0.95 in diploid genotypes (Hui et al. 2020). Among common variants in the HRC panel, 853,159 overlap with the 1240K SNPs panel.

1000 Genomes Project data were used to construct the reference haplotypes. We kept the standard parameters in hapROH, which had been optimised for 1240K aDNA genotype data:

> e_model=’haploid’, post_model=’Standard’, random_allele=True, roh_in=1, roh_out=20, roh_jump=300, e_rate=0.01, e_rate_ref=0.0, cutoff_post=0.999, max_gap=0, roh_min_l=0.01

### LSAI sharing and individual connectedness inference

LSAI segments and kinship coefficients were estimated from merged plink files of 61 imputed ancient genomes, 503 Europeans from the 1000 Genome Project and UK Biobank data with IBIS version 1.20.9 using different minimum shared segment length (-min_L) threshold – 4 cM for population genetic inference and 5 and 7 cM for kinship analyses - together with -maxDist 0.1 and -mt 300 parameters. In total, 269,319 binary SNPs with MAF >0.05 were used. Probabilities of IBD sharing among groups were estimated as in Kivisild et al. 2021.

### Phenotype prediction

For 39 out of the 41 HIrisPlex-S set of SNPs we selected 2 Mb around the informative variants, merging the regions on the same chromosome, with the exception of the variants on chromosome 15, which have been analysed in two different regions since the distance between the two nearest SNPs was about 20 Mb. We selected 10 regions from 9 autosomes, spanning from about 1.5 Mb to 6 Mb. For the other phenotypic informative markers (diet, immunity and diseases), we selected 2 Mb around each variant and merged the overlapping region, for a total of 48 regions from 17 autosomes and the X chromosome.

We call the variants using ATLAS v0.9.0 (Link et al. 2017) task=call and method=MLE commands at positions with a minimum allele frequency (MAF) ≥ 0.1% in the reference panel, that has been selected according to the different components of the samples: 1) Europeans from 1000 Genomes (EUR) (1000 Genomes Project Consortium et al. 2015) for Mesolithic, Neolithic, Copper Age and Bronze Age samples; 2) UK10K individuals extracted from the Haplotype Reference Consortium (HRC) (McCarthy et al. 2016) (accessed at: http://www.haplotype-reference-consortium.org/) for Iron Age, Roman and Early Medieval individuals from Great Britain; 3) EUR plus the MANOLIS (EUR-MNL) set from Greece and Crete extracted from the HRC (McCarthy et al. 2016) for the Imperial and Later Romans from (Antonio et al. 2019) and already analysed for the same phenotypic variants in (Saupe et al. 2021). For both the unpublished and published set of samples described above, the variant rs34536443, a risk factor for tuberculosis with fluctuating frequency in Europe over the last 2,000 years (Kerner et al. 2021), has been analysed here for the first time. After calling the variants separately for each sample, we merged them in one VCF file per region. We used the merged VCFs as input for the first step of our imputation pipeline (Hui et al. 2020) (genotype likelihood update), performed with Beagle 4.1 -gl command (Browning and Browning 2016) using the same panels as before as reference. We then discarded the variants with a genotype probability (GP) less than 0.99 and imputed the missing genotype with the -gt command of Beagle 5.0 (Browning et al. 2018) using the HRC as a reference panel for all groups of samples. We then discarded the variants with a GP < 0.99 and used the remaining SNPs to perform the phenotype prediction. Two markers of the HIrisPlex-S set, namely the rs312262906 indel and the rare (MAF=0 in the HRC) rs201326893 SNP, were not analysed because of the difficulties in the imputation of such variants. Results are reported in Data S6.

We performed this analysis on new and previously published ancient samples with a coverage ≥ 0.04x. We then grouped the individuals in different cohorts depending on both time and space. First, we grouped the ancient individuals from the British Islands in five groups from the Mesolithic to the Early Medieval period and compared them with the modern GBR. We compared the groups with a sample size higher or equal to 5 performing an ANOVA test and, for the significant variants, we performed a Tukey test to identify the significantly different pairs of groups (Data S6B,C). Using the same approach, we also analysed the difference within Iron Age and Roman Britain by creating 8 local groups and comparing them with ancient Roman Italians, discarding the groups with a sample size less than 5. For both comparisons, we used a Bonferroni’s correction on an alpha value of 0.01 for the number of tested SNPs to set the significance threshold. We performed our phenotype analysis in a set of 285 ancient individuals, composed of the 43 new individuals reported here for the first time, 231 ancient people from the British Islands and 11 ancient Italians from the literature and already analysed for phenotypes in (Saupe et al. 2021). Sample-by-sample phenotype prediction and genotype at the selected phenotype informative SNPs, reported as the number of effective alleles (0, 1 or 2) are shown in Data S6.

### Male-biassed kinship in Prehistoric Britain and Ireland

Information regarding archaeological sites and genomes were pulled from the annotation file of the Allen Ancient DNA Resource (AADR) (Anon) and the supplementary materials of Olalde et al. 2018, Brace et al. 2019, Cassidy et al. 2016, Cassidy et al. 2020, Sanchez-Quinto et al. 2019 and Scheib et al. 2019.

For each neolithic site the total number of estimated individuals, the number of genomes retrieved and number of genomes tested for kinship were noted.

## Acknowledgements

We thank the support of the Cambridge Archaeological Unit, the other members of the After the Plague project (Piers Mitchell, Bram Mulder, and Jay Stock), the Estonian Biocentre and aDNA group for their help and expertise, and Stephen Hoper and Paula Reimer and the 14Chrono Centre at Queen’s University Belfast for assistance with the radiocarbon dates. This work was supported by the Wellcome Trust (Award no. 2000368/Z/15/Z) and St John’s College, Cambridge (J.E.R., S.A.I, C.C., A.R., T.O.C., C.L.S.); the Estonian Research Council grant PUT (PRG243) (A.S., M.M., C.L.S); and the European Union through the European Regional Development Fund (Project No. 2014-2020.4.01.16-0030) (C.L.S., M.M.); the European Regional Development Fund (Project No. 2014-2020.4.01.15-0012) (M.M.). A.R acknowledges support from the British Archaeological Association.

## Authors’ contributions

Conceptualization - C.L.S., J.E.R, and T.K.

Data curation - C.L.S., R.H.

Formal Analysis - C.L.S., R.H., T.K., S.B., G.R., E.D.’A and A.K.R.

Funding acquisition - J.E.R, T.K., M.M.

Investigation - C.L.S., S.J.G., A.S., A.K.R., S.S.

Methodology - C.L.S., T.K.

Project administration - C.L.S., T.K., J.E.R.

Resources - C.C., S.A.I., C.E., B.N., R.W., T.B., C.D., S.J.H.

Software - R.H., E.D.’A., S.B.

Supervision - C.L.S., T.K., T.O.C.

Validation - C.L.S., T.K., R.H., E.D.’A.

Visualisation - T.K., S.B., C.L.S.

Writing – original draft - C.L.S., A.R.

Writing – review & editing - All authors

## Availability of data and material

The accession number for the DNA sequences reported in this paper is ENA: under accession number PRJEB52707 (http://www.ebi.ac.uk/ena/data/view/PRJEB52707). The data are also available through the data depository of the EBC (http://www.ebc.ee/free_data).

## Supplementary Data

**Data S1. Sequencing Summary**

**Data S2. Comparative genomes**

**Data S3. F statistics**

**Data S4. IBD Sharing**

**Data S5. Runs of Homozygosity**

**Data S6. Phenotype prediction**

**Data S7. Isotope data**

